# Microgravity enhances the phenotype of *Arabidopsis zigzag-1* and reduces the Wortmannin-induced vacuole fusion in root cells

**DOI:** 10.1101/2022.04.01.486597

**Authors:** Mengying Wang, Katherine Danz, Vanessa Ly, Marcela Rojas-Pierce

## Abstract

The spaceflight environment of the International Space Station poses a multitude of stresses on plant growth including reduced gravity. Plants exposed to microgravity and other conditions on the ISS display root skewing, changes in gene expression and protein abundance that may result in changes in cell wall composition, antioxidant accumulation and modification of growth anisotropy. Systematic studies that address the effects of microgravity on cellular organelles are lacking but altered numbers and sizes of vacuoles have been detected in previous flights. The prominent size of plant vacuoles makes them ideal models to study organelle dynamics in space. Here, we used *Arabidopsis zigzag1* (*zig-1*) as a sensitized genotype to study the effect of microgravity on plant vacuole fusion. Wortmannin was used to induce vacuole fusion in seedlings and a formaldehyde-based fixation protocol was developed to visualize plant vacuole morphology after sample return, using confocal microscopy. Our results indicate that microgravity enhances the *zig-1* phenotype by reducing hypocotyl growth and vacuole fusion in some cells. This study demonstrates the feasibility of chemical inhibitor treatments for plant cell biology experiments in space.

## Introduction

Growing healthy plants in space is a major goal towards the viability of long-term space missions as plants can provide food for astronauts and benefit mental health. The space environment includes multiple stresses for plant growth including different levels of gravity, radiation, decreased nutrient delivery and water recycling ^1,2^. The International Space Station provides an ideal experimental setting for the study of plant growth and development under space conditions and many experiments have been conducted with plants on the ISS ^1,3^. Reduced gravity on the ISS results in disoriented plant growth unless light is provided to activate directional growth from phototropism. Effects of the spaceflight environment on the ISS include reduced cell elongation ^4^ or altered cell division ^5^ in roots. At the cellular level, the microtubule and actin cytoskeletons as well as the cell wall are altered under microgravity ^6–8^. Other cellular organelles are altered under microgravity conditions include mitochondria, plastids and ER ^9^. Interestingly, vacuole numbers increased in root apical cells for *Arabidopsis* seedlings grown on the ISS ^10^, and vacuole volume increases in certain cells in microgravity ^11,12^. Moreover, proteins involved in intracellular membrane trafficking were found to show altered abundance in microgravity ^13–15^. Those results suggest vacuolar development as well as trafficking are disrupted in microgravity, but overall, there is a large gap in documenting the effect of spaceflight on the organization of cellular organelles ^16^.

The plant vacuole is a large essential organelle in plant cells and is important for cell elongation, recycling, and storage, among many other functions ^17–19^. Vacuole enlargement is concomitant with cell growth along the developmental gradient represented by the longitudinal axis of the root. Vacuole enlargement along this gradient requires homotypic membrane fusion, which is mediated by two conserved complexes, the homotypic vacuole protein sorting (HOPS) and soluble N-ethylmaleimide-sensitive factor attachment protein receptor (SNARE) ^20^. Both HOPS and the vacuolar SNARE complexes have vacuole or pre-vacuole-specific subunits that provide specificity to membrane fusion. The vacuolar SNARE complex is formed by SYP22, VTI11, SYP51, and either VAMP727 ^21^ or VAMP713/VAMP711 ^22–24^. A loss of function allele of VTI11, *zigzag1* (*zig-1*), has fragmented vacuoles and showed abnormal shoot gravitropism ^25–27^. The *zig-1* mutant also has short hypocotyls, small and wrinkled rosette leaves and a zigzag-shaped stem ^25,26^. Wortmannin (WM), a chemical inhibitor of phosphoinositide 3-kinase (PI3K), can induce vacuole fusion in *zig-1* and partially restore its agravitropic shoot phenotype ^27,28^. WM-induced vacuole fusion in *zig-1* is fast and almost complete vacuole fusion can be observed in roots within two hours ^27,28^. Given the large size of the vacuole in plant cells and the efficiency of WM at inducing its fusion, this treatment represents a unique assay for the study of organelle fusion in plants in space.

Plant specimens have been sent to space for fundamental biology research since the 1960s ^1,3,29^ and this resulted in the design of specific hardware for optimal growth ^3,30^. The Biological Research in Canister-Petri Dish Fixation Unit (BRIC-PDFU) is a fully contained hardware that has been used extensively with small plant seedlings ^10,31–33^ and it is suitable for chemical treatments in space. Previous experiments involved storing liquid media or fixative solution inside the chamber of the PDFU for microbial culture or plant tissue fixation ^34–36^. Cell biology research on the ISS requires sample fixation to preserve cell morphology. Chemical fixation in the ISS has been accomplished with non-coagulant agents such as formaldehyde and glutaraldehyde^10^ but the latter has high levels of autofluorescence ^37^. Direct fluorescence microscopy of fluorescent proteins from fixed tissues ^38,39^ is an effective approach to extract quantitative information of cell structure at high samples size, but the viability of fluorescence microscopy of tissues fixed in space has not been demonstrated given the challenges imposed by the BRIC hardware.

Here, we took advantage of the WM-induced vacuole fusion in *zig-1* to study the effect of microgravity on plant vacuole fusion. Formaldehyde-based tissue fixation in the BRIC-PDFU hardware was used and GFP-tagged vacuoles were imaged post-flight by confocal microscopy. We detected a mild but significant decrease in plant vacuole fusion in *zig-1* that highlights possible effects of microgravity or other spaceflight stresses on vacuole biogenesis and membrane fusion. This experiment also demonstrates the feasibility of chemical treatment assays on the ISS which may contribute to future cell biology research in this microgravity environment.

## Results

### Optimization of BRIC-24 workflow

The goal of BRIC-24 was to test whether vacuole fusion is affected by the microgravity environment of the ISS. A previously characterized vacuole fusion assay ^27,28^ was implemented with the BRIC-PDFU hardware. This assay takes advantage of the accumulation of vacuoles in the *zig-1* mutant and the rapid induction of vacuole fusion in this mutant upon treatment with Wortmannin (WM) ^28^. Visualization of organelle morphology is accomplished by confocal microscopy of a GFP-tagged vacuolar marker (GFP-TIP2;1) ^40^ and a mCherry-tagged ER marker (HDEL-mCherry) ^41^ previously characterized. Figure 1 describes the BRIC-24 workflow as it was completed as part of the SpaceX-22 mission in July of 2021. Thirty sterile seeds from WT (Columbia-0) or *zig-1* were plated directly on *Arabidopsis* Growth Media (AGM) plates before integration into the BRIC-PDFU hardware at Kennedy Space Center. From this point, plates were maintained cold and in the dark. Each PDFU was separately loaded with a treatment solution (WM or DMSO control in AGM) as well as fixative solution taking advantage of the dual chamber of the PDFU hardware. The fixative solution was infused with N2 gas to enhance its long term stability as described ^42^. Four BRICs containing a total of 22 PDFUs and 2 HOBO temperature recorders were delivered to NASA’s cold stowage facility prior to launch (Supplementary Table 1). Two days after arriving at the ISS, BRICs were unpacked and allowed to come up to the temperature of the ISS environment to promote germination and growth. WM or DMSO control solutions were applied at day 4 of seedling growth followed by fixative solution 2 hours later. BRICs were returned to a 4°C storage compartment until the return flight 4 weeks later, de-integration and confocal analysis at North Carolina State University.

**Figure 1.**
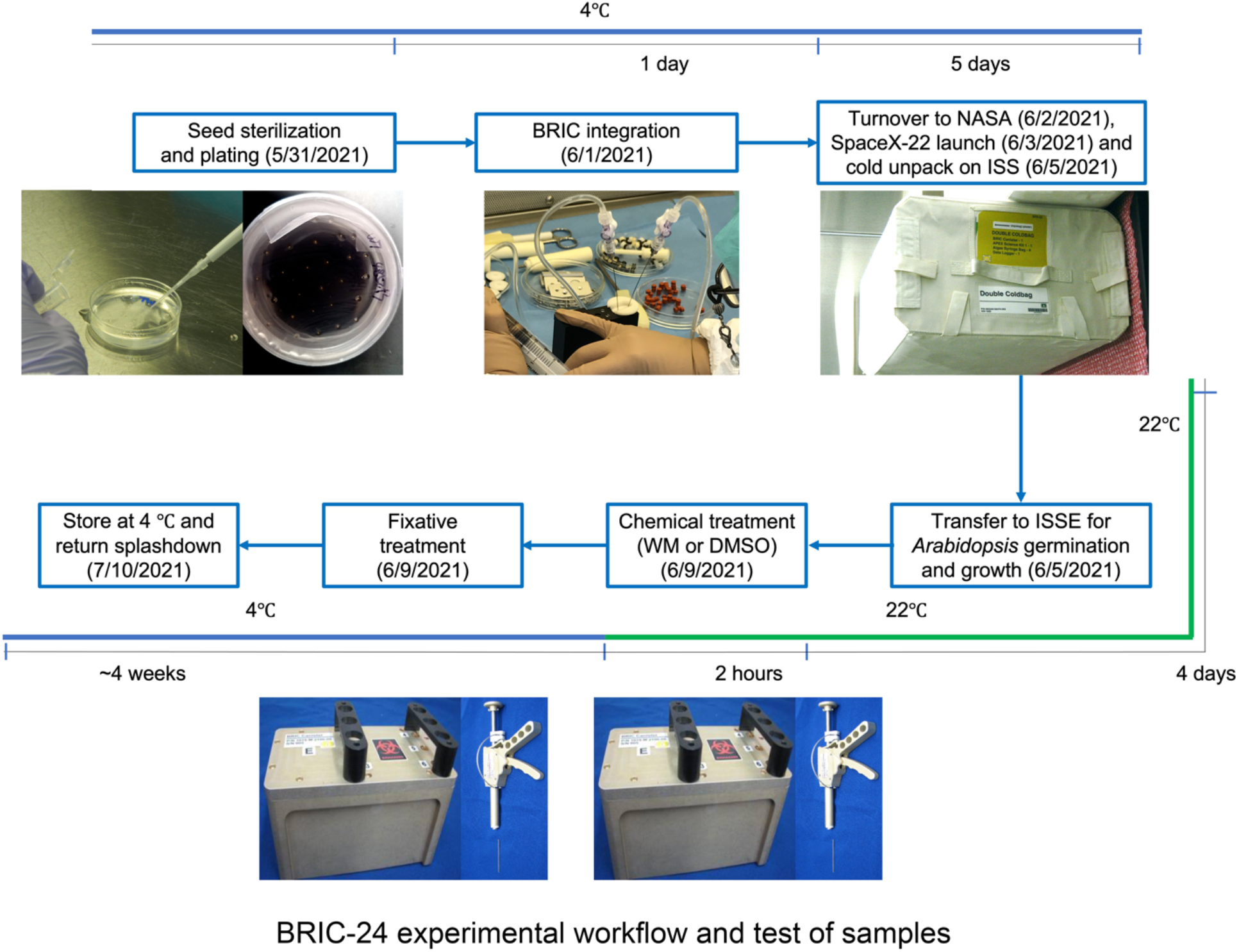
A schematic overview of the BRIC-24 experimental workflow. Sterilized *Arabidopsis* seeds (WT and *zig-1* with GFP-TIP2;1) were plated onto AGM media and stored at 4°C before BRIC-PDFU integration. Each petri dish contained 30 seeds. Treatment solution (33 μm WM or DMSO buffer) and fixative were injected into the dual chamber of the PDFU. Plates were placed into each PDFUs and PDFUs were integrated into BRICs. Four BRICs were then stored in a cold bag for the flight. After 4 days of growth on the ISS, WM solution or DMSO control were applied to the seedlings. The treatment lasted for 2 hours. The second actuation was conducted by applying fixative solution to the seedlings. BRICs were then kept at 4°C and returned to NASA after splash down. Confocal imaging and data analysis were conducted at NCSU.

BRIC-24 protocols that fit the constraints of the flight schedule and hardware required considerations for storage of treatment solutions in the BRIC hardware before actuation and the preservation of treated and fixed seedlings for up to 6 weeks before imaging. First, we confirmed that 30 μM working dilutions of WM in liquid AGM buffer are still active after 2 weeks of storage at 4°C (Supplementary Fig. 1). Furthermore, we determined that the formaldehyde-based fixative solution was effective after 2 weeks storage (Supplementary Fig. 2). These results demonstrated the feasibility of the BRIC-24 workflow.

Transfer of the treatment and fixative solutions from the liquid chambers of the PDFUs and into the plants is carried out by astronauts in space, but this device does not allow for the removal of the first solution. Therefore, the fixative solution would always mix with the previously applied treatment solution. We then tested the effectiveness of a variety of fixative buffers and PFA concentrations when mixed 1:1 with liquid AGM by visualization the GFP-labeled tonoplast and the mCherry-labeled ER of fixed samples (Supplementary Fig. 3). A 4% working concentration of PFA was used as the fixative agent ^38,43^. When PFA was directly diluted in liquid AGM, the intensity of GFP fluorescence was weak, and the morphology of vacuole and ER were blurry compared to live-seedling controls (Supplementary Fig. 3 B, C). As most PFA fixation solutions are made in PBS and many variations exist, we tested different PBS concentrations and mixed them 1:1 with AGM. Fixative solutions in full (1x) PBS buffer as well as 0.5x PBS yielded plants with high intensity of GFP, however, only the PFA solution in 0.5x PBS showed the best preservation of vacuole morphology (Supplementary Fig. 3 D, E). In the case of the KPBS buffer, a variation of PBS that contains K^+^ as an alkali metal cation in addition to Na^+^ ion, vacuole morphology was acceptable, but GFP intensity was diminished (Supplementary Fig. 3 G, H). Weak GFP signal was also detected for samples fixed with fixative in MTSB buffer, a buffer typically used to preserve cytoskeleton in plants ^44^, (Supplementary Fig. 3 I, J). PFA solutions in PPB and Sorensen’s buffer were ineffective in preserving vacuole morphology (Supplementary Fig. 3 K, L). Preservation of ER morphology as reported by the mCherry-HDEL marker was even more challenging. Most fixative solutions resulted in the accumulation of ER aggregates and overall loss of ER structure, except for a PFA solution in 0.25x KPBS (Supplementary Fig. 3H). This solution, however, was not optimal for visualization of vacuoles. As vacuole fusion was the primary target for this research, we selected the 4% PFA solution in 0.5x PBS (Supplementary Fig. 3 E) as the final fixative concentration for the flight experiment and did not analyze ER morphology as part of BRIC-24.

One major concern for doing plant cell biology experiments in space is that fixed samples must be stored on the ISS for several weeks before splash-down. We therefore tested whether 4 weeks of storage would affect the intensity of GFP or the overall preservation of vacuole morphology of fixed samples. Several experiments confirmed that long term storage of these samples would not significantly affect their quality of preservation (Supplementary Fig. 4) further supporting the feasibility of BRIC-24.

### Ground tests demonstrated feasibility of the vacuole fusion assay using the BRIC-PDFU

Science (SVT) and Experiment (EVT) Verification Tests were completed at Kennedy Space Center on March and September 2020, respectively, to determine the feasibility of the experimental workflow using the BRIC-PDFU hardware. The PDFU is an enclosed environment where plants are exposed to dark conditions and where plant growth or the effectiveness of chemical treatments cannot be assessed in real time. It was important to test whether WM treatment of WT and *zig-1* inside the PDFU was equivalent to treatments regularly done on plastic dishes. Both SVT and EVT were used to optimize growth conditions in the PDFU and to optimize the WM treatment and fixation. Plants were grown for 4 days during SVT, and they were treated with 33 μM WM for 1 h before fixation in 4% formaldehyde in 0.25x KPBS. The germination rate during SVT was not significantly different between genotypes (Supplementary Fig. 5A), but the hypocotyl length of WT was longer than that of *zig-1* (Supplementary Fig. 5B) as previously reported ^25^. Fixed seedlings were imaged by confocal microscopy to visualize the vacuoles in the epidermis and cortex from the root transition and elongation zones (Supplementary Fig. 5C). Visualization of vacuoles after PFA fixation in KPBS buffer results in weak GFP fluorescence. As expected, a single vacuole was observed in most cells of WT roots regardless of chemical treatment. Multiple vacuoles were detected in *zig-1* roots in the control treatment and the WM treatment resulted in fewer vacuoles per cell when compared to the control. Most cells did not complete vacuole fusion after 1 h under these conditions and this was unexpected ^27,28^. This result could be due to hardware limitations or suboptimal fixation, and it prompted an increase in the incubation time of the WM in subsequent experiments.

A final EVT was conducted to demonstrate the feasibility of BRIC-24 before flight. Plants were grown for 3 days and treated with 33 μM WM for 2 h before fixation in 4% formaldehyde in 0.5x PBS to preserve GFP fluorescence (Fig. 2; Supplementary Fig. 5 D, E). Again, no significant differences in seed germination were detected between genotypes but *zig-1* hypocotyls were shorter than the WT (Fig. 2A; Supplementary Fig. 5 D, E). Root cortical cells of the root transition zone were imaged for vacuole morphology (Fig. 2 B, C) and most WT cells contained only one vacuole regardless of treatment. Multiple vacuoles per cell were observed in the *zig-1* control, and WM treatment resulted in a significant reduction in the number of vacuoles per cell in the mutant. These results suggest that chemical inhibitor treatments in the BRIC-PDFU are feasible, albeit adjustments must be made to accommodate changes due to the limitations of the hardware.

**Figure 2.**
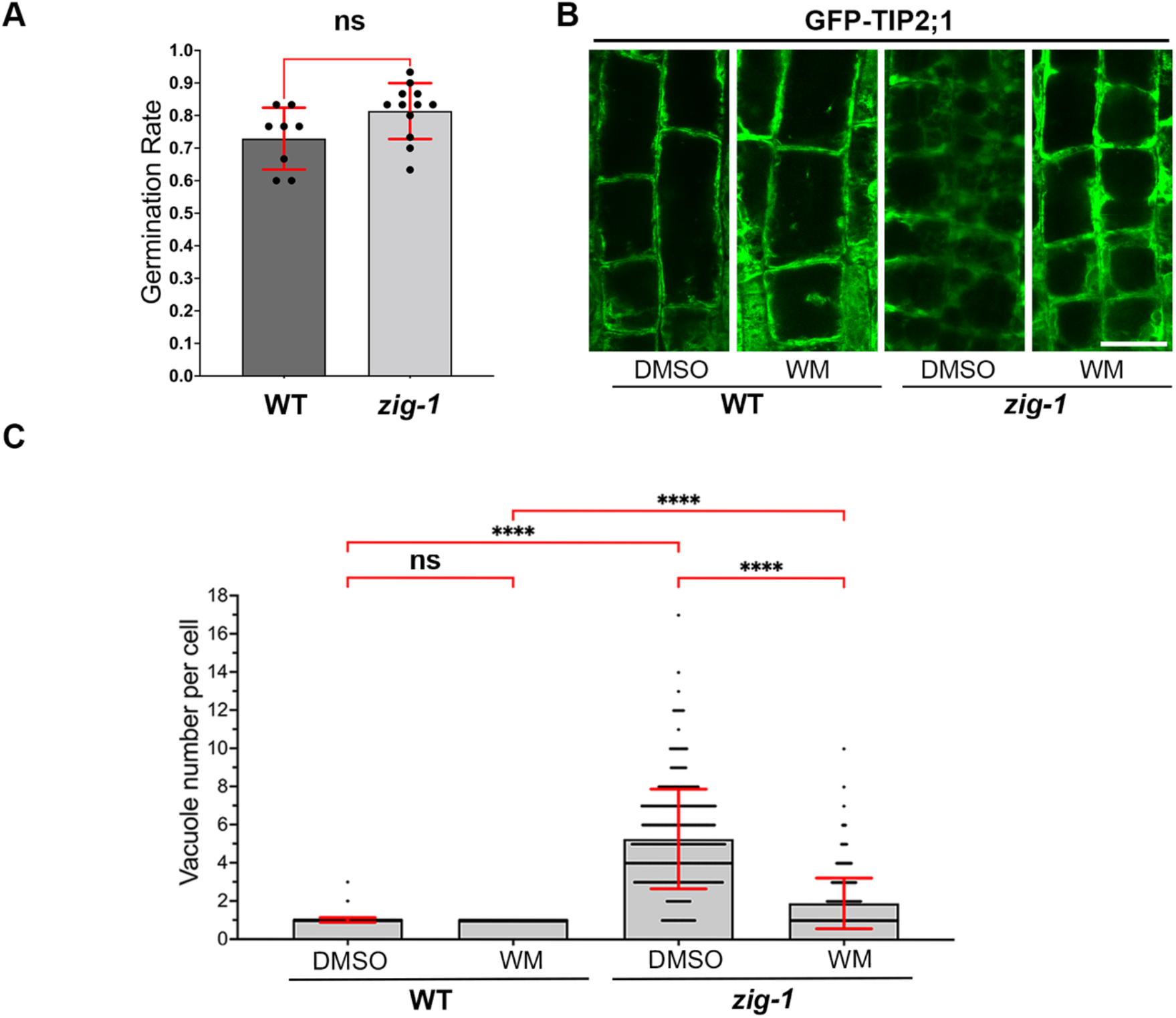
The WM-induced vacuole fusion assay can be carried out in the BRIC-PDFU. **(A)** Germination rate of WT and *zig-1* seeds inside BRIC-PDFU during EVT. No significant difference in germination rate between two genotypes was detected after removing the two WT outliners (30% and 50%). (Unpaired t test, ns: p>0.05; n represents plate number, n>=8) **(B)** Fixed root tip cortex cell vacuole morphology of WT and *zig-1* (GFP-TIP2;1). Three-day-old WT and *zig-1* seedlings were treated with DMSO buffer control or WM solution for 2 hours and then seedlings were fixed and kept at 4°C before confocal imaging. **(C)** Quantification of vacuole number in WT and *zig-1* after chemical treatment. Cells from the cortex transition zone of the root tip were analyzed. (Two-way ANOVA, Tukey’s multiple comparison test, **** p<0.0001; n represents cell number under analysis for each genotype with each treatment, n>365) Scale bar: 20μm

Even though the feasibility of BRIC-24 was demonstrated during EVT, the overall quality of GFP signal, seedling preservation and WM effectiveness was not on par with similar experiments outside the BRIC-PDFU hardware. This may result from the lack of precision when loading solutions into the dual chamber. We then tested whether deviations from the 1:1 ratio between the WM (or DMSO) solution in AGM and the fixative could affect the quality of cell fixation. Dark grown *zig-1* seedlings were fixed in different mixtures of liquid AGM and fixative solution that mimic unequal loading of each solution into the dual chamber. The best cell morphology and GFP signal preservation were obtained when those solutions were mixed with 1:1 ratio. Higher concentrations of fixative induced cell damage while lower concentrations of fixative resulted in lower GFP intensity (Supplementary Fig. 6). These results explain some suboptimal cell preservation and GFP fluorescence during SVT and EVT (Supplementary Fig. 7) and must be considered for future flight experiments using the BRIC-PDFU. Nonetheless, sufficient seedlings with good preservation and bright GFP fluorescence were recovered from EVT to demonstrate flight readiness of BRIC-24.

### *zig-1* showed normal germination but altered hypocotyl growth on the ISS

BRIC-24 was flown with the SpaceX-22 mission (Fig. 1). Identical experiments with 4 BRICs containing 22 PDFUs were completed on the ISS (flight) and at Kennedy Space Center (ground control). Due to a power failure, the ground control experiment ran 5 weeks after the flight. We analyzed the germination rate, seedling hypocotyl length of WT and *zig-1* from flight assay and compared the data to ground control. Seed germination was not affected by the microgravity environment of the ISS in either genotype (Fig. 3A). Like SVT and EVT experiments, WT hypocotyls were 7.35% and 31.67% longer than those of *zig-1* in ground control and flight respectively. However, the ISS environment resulted in 8.46% longer hypocotyls on the WT compared to ground control while the effect in *zig-1* was the opposite, where *zig-1* plants grown on the ISS were 11.57% shorter than the ground control (Fig. 3 B, C). These results suggest that seed germination of *zig-1* is not affected by microgravity, but its growth may be inhibited by the stresses imposed by the ISS environment.

**Figure 3.**
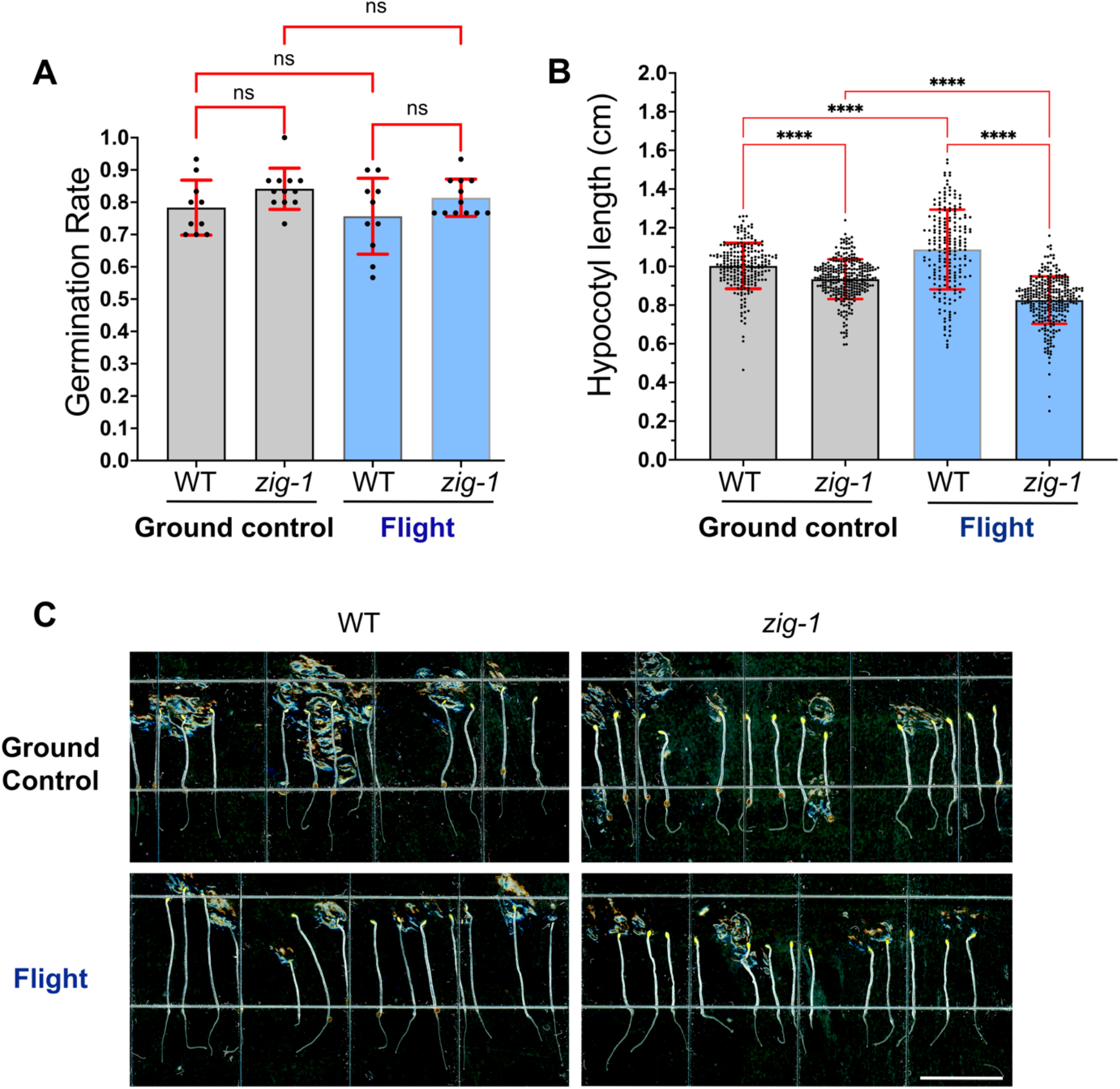
WT and *zig-1* in BRIC-PDFU can germinate and grow in microgravity environment as on earth. **(A)** Germination rate of WT and *zig-1* grown in the BRIC-PDFU from ground control or flight. No differences were detected in the germination rate between genotypes or treatments. (Two-way ANOVA, Tukey’s multiple comparison test, ns: p>0.1234, n represents plate number per genotype per treatment, n>10) **(B)** Hypocotyl length of WT and *zig-1* grown in the BRIC-PDFU from ground control and flight. WT showed longer hypocotyl than *zig-1* both for ground control and flight. Flight assay promoted the hypocotyl length of WT while inhibited *zig-1’s* hypocotyl length. (Two-way ANOVA, Tukey’s multiple comparison test, **** p<0.0001, n represents seedling number per genotype per treatment, n>200) **(C)** Representative seedlings of the WT and *zig-1* from either ground control or flight after de-integration of the BRIC-PDFU. Fixed seedlings were taken out of fixative solution and arranged on plate. Scale bar: 1cm

### WM-induced vacuole fusion may be delayed in microgravity

The primary goal for this research was to test whether WM can induce vacuole fusion of *zig-1* in the microgravity environment of the ISS. For this purpose, all seedlings collected from BRIC-PDFUs of flight assay and ground control were imaged by confocal microscopy for vacuole number analysis. Epidermal and cortical cells within the transition and elongation zones of the root tip were analyzed as vacuole morphology in these cells is developmentally controlled ^45,46^. WT cells had only one vacuole with a few exceptions in the transition zone, regardless of chemical treatment or flight environment (Fig. 4). This suggests that the microgravity environment on the ISS does not induce major delays on vacuole biogenesis in the WT. *zig-1* seedlings showed the typical fragmented vacuole phenotype in both the flight assay and the ground control when the DMSO control treatment was applied (Fig. 4). While no differences were observed between flight and ground control in the *zig-1* transition zone of roots treated with DMSO (Fig. 4 B, F), a small but significant increase in vacuole number was detected in the elongation zone of flight samples in *zig-1* (Fig. 4D, H). For example, the average vacuole number per cell was 3.27 for ground control and 3.80 for flight assay in the epidermis of the elongation zone. Therefore, *zig-1* may have an enhanced *zig-1* phenotype in the microgravity environment of the ISS.

**Figure 4.**
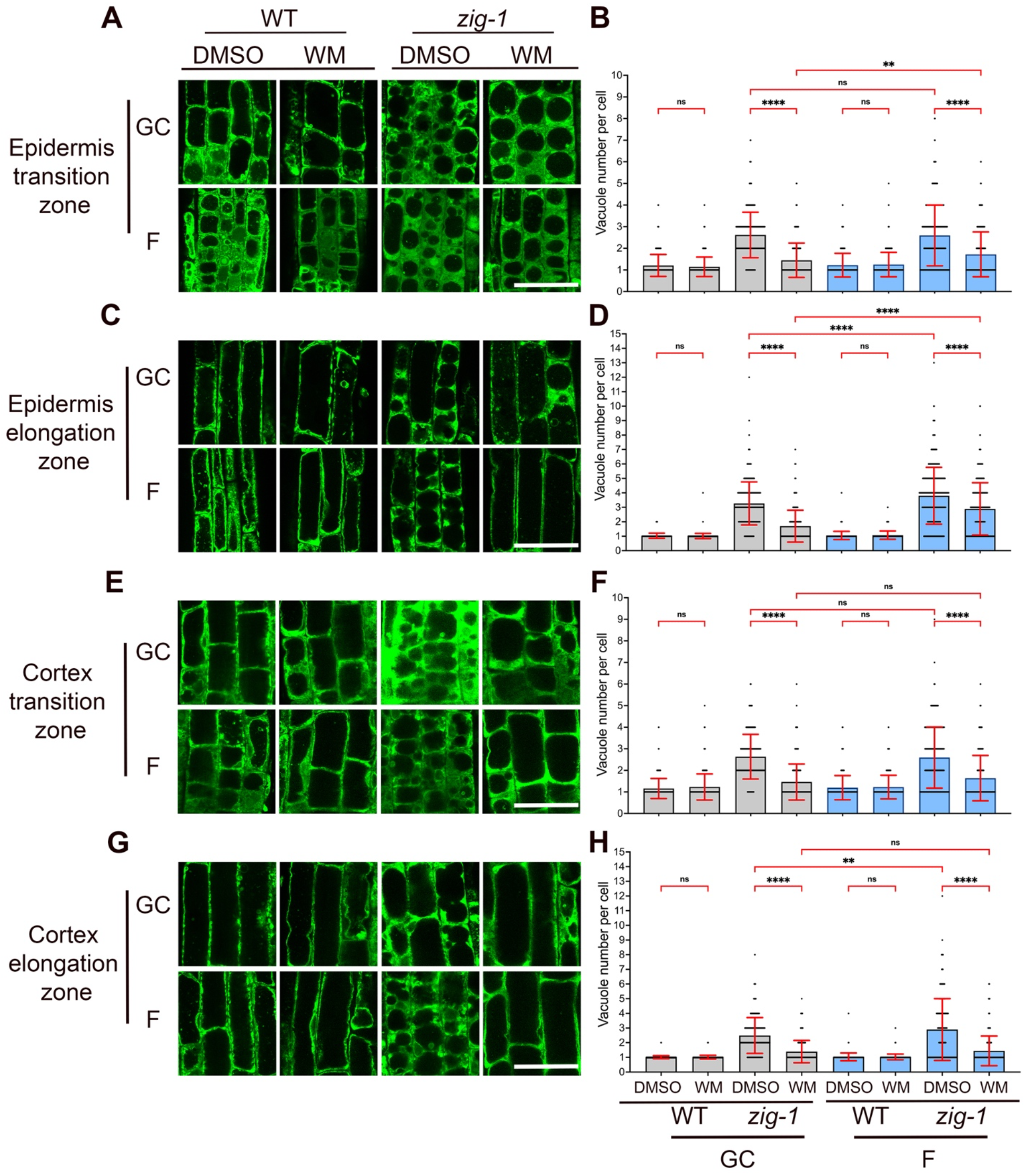
Quantification of WM-induced vacuole fusion. Vacuole morphology (**A, C, E, G**) and vacuole number (**B, D, F, H**) of WT and *zig1* plants expressing GFP-TIP2;1. Seedlings were grown in ground control or the ISS and treated with DMSO buffer (control) or WM solution. Cells were imaged at the root tip epidermis transition zone (**A, B**), epidermis elongation zone (**C, D**), cortex transition zone (**E, F**) or cortex elongation zone (**G, H**). Four-day-old seedlings were treated with DMSO buffer control or WM solution for 2 hours and then seedlings were fixed and kept in 4°C before confocal imaging. (Three-way ANOVA, Tukey’s multiple comparison test, **** p<0.0001; **p<0.0021; n represents cell number under analysis per genotype per treatment, n>155) Scale bar: 40 μm

Vacuole numbers in WM-treated plants was used as a proxy for vacuole fusion events (Fig. 4). As expected, the vacuole number per cell in *zig-1* treated with WM was smaller than the DMSO control for both flight and ground control samples and in all cells analyzed. This suggests that the WM treatment was effective at inducing vacuole fusion inside the PDFU. Like the EVT experiment, WM did not induce complete vacuole fusion in either flight or ground control as *zig-1* cells containing more than one vacuole were observed in all tissues. However, a higher number of vacuoles was observed in the epidermis of WM-treated *zig-1* grown on the ISS when compared to the ground (Fig. 4 A-D, Supplementary Table 2). These results suggest a small delay in *zig-1* vacuole fusion during flight. This hypothesis is supported by the smaller difference in the average vacuole number between DMSO and WM treatments in the epidermis transition zone for the flight samples (0.88) when compared to ground control (1.17). Similar results were found in the epidermis elongation zone, with these differences being 0.91 (flight) and 1.57 (ground), respectively. This sensitivity to the flight environment was not detected in the cortical cells where the difference between WM and DMSO vacuole numbers were similar in the two environments. These results overall suggest a cell-type specific response to the stress conditions of the flight environment that resulted in small decreases in vacuole fusion in the epidermis.

### Amyloplasts were associated with tonoplast in microgravity

Vacuoles are tightly associated with other organelles and vacuole-amyloplasts associations might be involved in gravity sensing ^28^. We took advantage of the strong GFP fluorescence of fixed BRIC-24 samples and the vacuole phenotype of *zig-1* to test whether microgravity alters the association of vacuoles with amyloplasts. Statocytes from WT and *zig-1* seedlings that were treated with the DMSO buffer were imaged by confocal microscopy (Fig. 5; Supplementary Fig. 10). Amyloplasts in hypocotyl endodermal cells were found closely associated with tonoplast both for flight sample and ground control and in both genotypes, and the tonoplast membrane surrounded all plastids (Fig. 5). As expected, WT amyloplasts were found at the bottom of the cell in the ground control samples while they were distributed throughout the cell in the ISS samples. Like previous results ^28^, amyloplasts appeared trapped between fragmented vacuoles in *zig-1* for both flight and ground control. Unfortunately, we could not analyze root statocytes in a similar manner due to low GFP fluorescence intensity in root columella cells (Supplementary Fig. 10). These results overall suggest that the microgravity environment of the ISS does not appear to alter the association of vacuoles and amyloplasts at least in hypocotyls.

**Figure 5.**
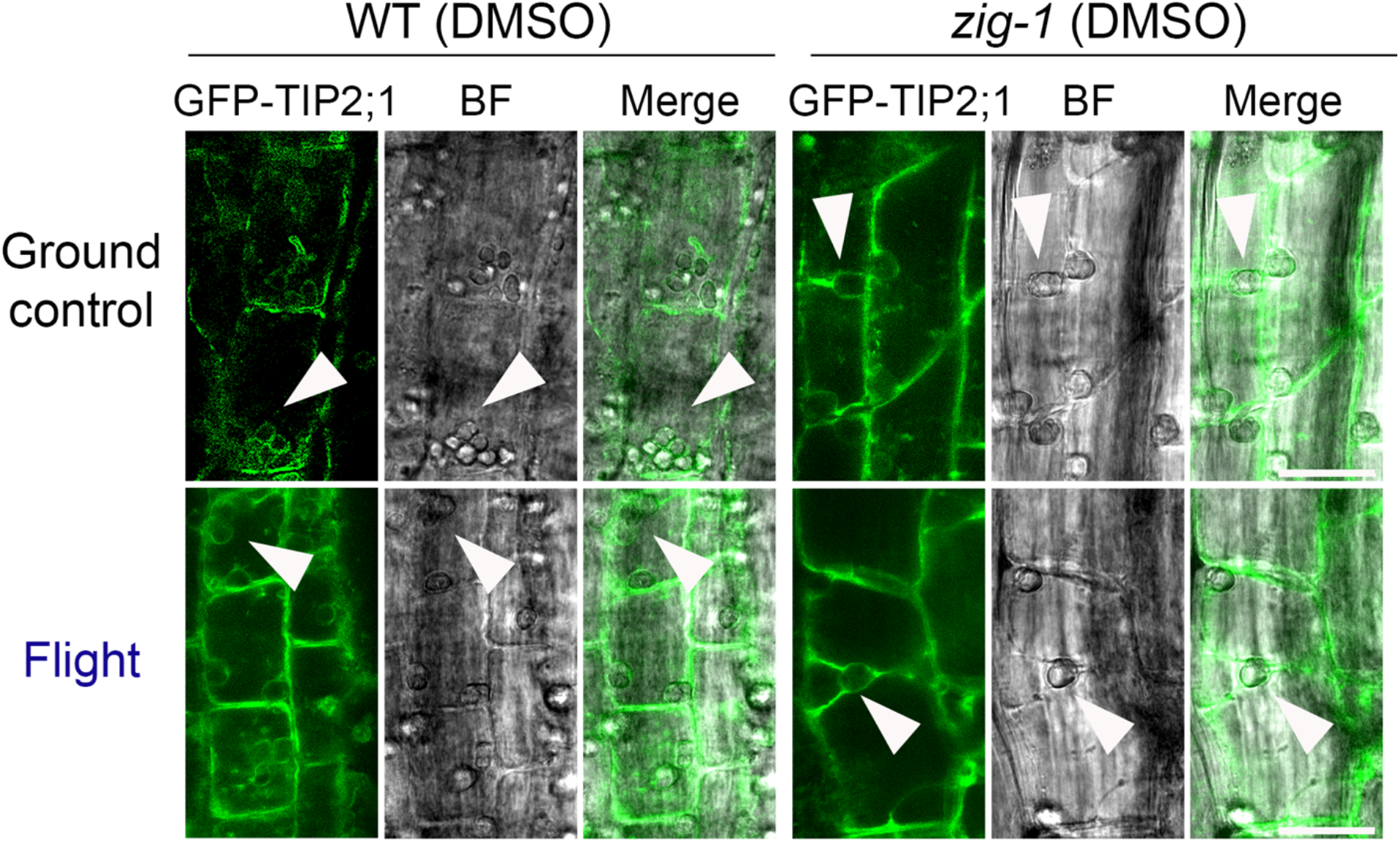
Visualization of the vacuole and amyloplast inside the hypocotyl endodermal cell. Amyloplasts (where arrows point at) were associated with tonoplast (labeled by GFP) in hypocotyl endodermis of WT and *zig1* in microgravity environment as on earth. Scale bar: 20μm

## Discussion

Here, we investigated the effect of microgravity on plant vacuole fusion and the association of vacuoles and plastids. The feasibility of chemical treatments in plants on the ISS was demonstrated using the BRIC-PDFU hardware and an optimized plant fixation protocol.

### The microgravity environment of the ISS enhances the phenotype of *zig-1*

WT seedlings had longer hypocotyls when grown on the ISS and this was consistent with previous experiments with seedlings grown in the dark ^47–49^. The increased elongation of hypocotyls in microgravity may result from increased cell wall extensibility and reduced accumulation of xyloglucans ^47^. In addition, an increase in transverse microtubules may also be involved in changing the rate of cell elongation ^49^. On the other hand, *zig-1* mutants showed shorter hypocotyls and more vacuoles per cell in microgravity-grown seedlings. This suggests an enhanced *zig-1* phenotype on the ISS. Given the role of VTI11 as a SNARE for protein trafficking ^50^, it is tempting to speculate that the microgravity environment may have a negative effect for endomembrane trafficking overall. Moreover, the enhanced phenotype of *zig-1* suggests that vacuole fusion is decreased in microgravity even though this can only be detected in this sensitized genotype. Cell elongation is in part dependent on the establishment of the large central vacuole ^18^ and the reduction in hypocotyl elongation in *zig-1* would therefore be consistent with a possible delay in vacuole fusion in that tissue. Given the lower expression of the GFP-TIP2;1 marker in hypocotyl cells, it was not possible to directly test this hypothesis during BRIC-24.

### The microgravity environment delays WM-induced vacuole fusion in *zig-1* epidermis

The WM fusion assay is an ideal tool for visualization of cellular membrane fusion in fixed tissues in space. We selected the cortex and epidermis from the transition and early elongation zones of the root because while the fragmented vacuole phenotype of *zig-1* is present in most cell types, the WM-induced fusion is best visualized in those root cells ^27,28^. Moreover, root cells undergoing the transition between meristem and elongation zones are well-defined stages of vacuole maturation ^46,51–53^. Overall, we found cells from the transition and early elongation zone to be better preserved than central elongation as well as the maturation zone. Given the large numbers of seedlings used, this allowed us to obtain quantitative information for this assay. The vacuole number analysis in epidermis and cortex indicated that WM was overall effective at inducing vacuole fusion of *zig-1* in microgravity inside the BRIC-PDFU. This result underscores the feasibility of chemical inhibitor treatments in plants grown on the ISS. These experiments, however, also exposed limitations rendered by the PDFU hardware as vacuole fusion was incomplete even in the ground control, unlike similar experiments outside the hardware where almost all the *zig-1* cells result in having a single vacuole. Given the sealed nature of the PDFU and the mode of actuation, contact of the chemical solution with all the seedlings on the plate may be incomplete and this would be exacerbated by the abnormal movement of liquids under microgravity. Nonetheless, a small but significant delay in WM-induced vacuole fusion was detected in the epidermis of flight-grown *zig-1*. This result, together with the increased vacuole number in this mutant under microgravity, suggests that microgravity may slow down membrane fusion events in plants but not sufficiently to inhibit growth or be detectable in the WT. Such effect of microgravity on vacuole fusion and the sensitized nature of the *zig-1* genotype may explain the reduced hypocotyl growth of *zig-1* under flight.

### Fixation of fluorescent protein-labeled organelles on the ISS is feasible

Previous cell biology experiments on the ISS relied on tissue fixation followed by electron microscopy ^10^. Fluorescence microscopes have been available at the ISS ^49,54^ which allowed the visualization of GFP tagged proteins in plants ^55^, but these have been limited to tissue expression patterns due to the complexity of high resolution imaging ^56^. Imaging of plant cells with high resolution requires high numerical aperture objectives with short working distances which require specimens be mounted on slides, something that would require a large amount of astronaut time. Therefore, tissue fixation and postflight microscopy is still the best option for cell biology when large sample numbers are needed. Electron microscopy of fixed samples provides the best cellular and subcellular resolution but is time consuming. We set out to develop effective protocols for the fixation of *Arabidopsis* seedlings carrying fluorescently tagged proteins such that we could take advantage of the capabilities and efficiency of confocal microscopy. We were able to obtain good quality fixations in both genotypes that allowed us to quantify vacuoles in sufficient seedlings, but it is important to emphasize that the fixation quality was still diminished compared to similar fixation protocols outside the BRIC-PDFU hardware even on earth.

The fixation protocol described here was optimized to follow a chemical treatment in AGM buffer. BRIC-PDFU actuations result in the mixture of fixative and chemical treatment solution as there is no mechanism to remove the first solution. Both the AGM and the fixative buffer contribute to the osmolarity of the final mixture and therefore affect fixation quality. Membrane vesiculation was observed in some fixed seedlings (Supplementary Fig. 3, 7), and further optimization of the fixation protocol may improve the post-flight imaging of stored fixed samples. Similar vesiculation phenomena has been reported in formaldehyde-fixed samples in other systems ^57^. A further challenge of the PDFU hardware that might impinge on fixation quality is the low precision of the volumes used to fill the dual chamber of PDFUs. Our study demonstrated that variations from the 1:1 ratio of AGM to fixative can dramatically alter the fixation quality of plant specimens. Future experiments using this hardware must ensure sufficient sample size to account for the loss of samples due to poor fixation. An improved dual chamber PDFU unit with more precise volume control would augment the use of this hardware in future flight experiments.

Tissue fixation protocols that are compatible with fluorescent proteins have been developed for plants ^38^,^39^,^43^,^58^. These protocols were developed to improve imaging depth within plant tissues by clearing cellular structures. Maintaining subcellular structure and fluorescent protein structure and intensity allow for imaging experiments that are incompatible with live-cell imaging such as those conducted in space. We found that the intensity of GFP and mCherry is retained in fixed seedlings even after long storage in the cold. However, GFP intensity was more sensitive than mCherry to buffer composition. Lower concentration of KPBS buffer reduces the intensity of GFP, suggesting the GFP stability can be altered by the concentration of certain salt. Unfortunately, the morphology of the mCherry-tagged ER was poorly preserved with most fixative solutions. Many aggregates of ER membranes were detected while the fluorescence intensity of the mCherry marker was intact. A similar phenomenon was reported in *Drosophila* cells carrying a BiP-Superfolder GFP-HDEL fusion that labels the ER. In those cells, chemical fixation disrupted the ER morphology and ER tubules appeared to fragment into discrete puncta ^59^. Preservation of ER structures by chemical fixation may require specific chemical compositions.

## Materials and methods

### *Arabidopsis* materials and growth conditions

*Arabidopsis* ecotype Columbia (Col-0) was used as WT control and *zig-1* mutant was previously described ^25^. Col-0 and *zig-1* seeds stably expressing HDEL-mCherry ^41^ and GFP-TIP2;1 ^40^ were used to visualize the ER and tonoplast under confocal microscopy.

Seeds were surface sterilized and sown on 0.5 X *Arabidopsis* growth medium (AGM, 0.5X Murashige and Skoog medium, 10g/L Gelrite, 1% (w/v) sucrose) in 6 cm diameter petri dishes. Each petri dish contained 30 seeds. Twenty-two Petri dishes were integrated into PDFUs and stored in 4 BRICs for SVT, EVT and flight. BRICs were incubated at 4°C for 5 days in the dark and transferred to 22°C growth chamber (SVT, EVT and ground control) or the ISS environment (flight assay) for 3-4 days (Fig. 1).

### Chemical treatments and fixation

Three to four-day-old dark-grown seedlings were used for chemical treatments and fixation. WM (SIGMA, cat# W3144) was diluted to 33 μM in AGM and an equal volume of DMSO (Thermo Scientific, D12345) diluted in AGM was used as the control. For lab testing, seedlings were incubated in WM working solution or DMSO buffer control for 1 or 2 hours in the dark. If used in the BRIC-PDFU, WM or DMSO were stored in one of the chambers in the PDFU and an actuator tool was used to push each solution onto the petri dish.

Optimization of the seedling fixation protocol involved testing different buffers for the PFA fixative. To mimic the mixing process inside the BRIC-PDFU, one volume of 0.5X AGM was mixed with one volume of 8% Paraformaldehyde (PFA, Electron Microscopy Sciences cat# 15700) in the corresponding buffer for a final concentration of 4% PFA. The final concentration for each buffer to test was 0.5X AGM, 0.5X or 0.25X KPBS ^60^, 0.5X or 1X BupH™ PBS (ThermoFisher Scientific cat# 28372), 0.5X or 1X microtubule stabilizing buffer (MTSB) ^44^, 1X Potassium phosphate buffer (PPB) (6.5mM KH_2_PO_4_, 93.5 mM K_2_HPO_4_) and 0.25X Sorenson’s buffer ^61^. Seedlings treated with each mixture were immediately stored at 4°C until confocal microscopy.

For SVT, EVT and flight assay, 16% PFA aqueous solutions were diluted to 8% PFA in 0.5X KPBS (SVT) or 1X BupH™ PBS (EVT, flight assay). The final PFA concentration after actuation in the PDFU was around 4% with 0.25X KPBS buffer (SVT) and 0.5X BupH™ PBS buffer (EVT, flight assay). Seedlings were then stored at 4°C until de-integration of BRIC-PDFU followed by confocal microscopy.

### Microscopy

Fixed seedlings were washed 3 times for 1 min in sterilized water before imaging. A Zeiss LSM880 confocal microscope was used for all the imaging. The excitation/emission wavelengths for GFP and mCherry were 488 nm / 490-561 nm and 561 nm / 570-676nm respectively. The C-Apochromat 40X/1.2 W Korr FCS M27 water-immersion objective was used. All seedlings were imaged at the root tip transition and elongation zones and in both epidermis and/or cortical cells. Some seedlings were imaged for root tip columella cells and hypocotyl endodermal cells.

### Image analysis

For SVT, EVT and the flight assay, seeds germination rate was calculated from the number of recovered seedlings in each PDFU after de-integration. To measure hypocotyl length, fixed seedlings were arranged on glass slides and imaged using digital cameras. Hypocotyl length was measured with Fiji ^62^.

To count the number of vacuoles per cell, images from different tissue and cell type was analyzed separately. Only cells with good preservation and strong GFP fluorescence intensity were counted. Three-way ANOVA analysis was applied to compare the cellular vacuole amount among different treatments. Statistical analysis and figure generation were conducted with GraphPad Prism version 9.2.0 for macOS, GraphPad Software, San Diego, California USA, www.graphpad.com.

## Acknowledgements

Special thanks to Dr. Ye Zhang from Kennedy Space Center, Gerard Newsham, Donald Houze, Susan Manning-Roach, Anne Marie Campbell form MEI Technologies for help with the SVT, EVT and Flight assays. Special thanks to astronaut Thomas Pesquet for conducting the BRIC-PDFU actuation during flight. This work was supported by a grant from the National Aeronautics and Space Administration (80NSSC19K0145 to M.R.P).

## Author contribution

MRP designed the experiment. MW, KD, VL and MRP carried out experiments. MW developed the fixation protocol, prepared plant samples, imaged samples, and analyzed data. MW and MRP wrote the manuscript.

## Data availability

All data of this study are available from the corresponding author upon request.

