## Supplementary Materials for "Microgravity enhances the phenotype of *Arabidopsis zigzag-1* and reduces the Wortmannin-induced vacuole fusion in root cells"

**Supplementary Table 1**. Genotypes and treatments used in the BRIC-PDFU.

| BRIC Name | Position of PDFU | Genotype | Treatment | Position of PDFU | Genotype | Treatment |
|---|---|---|---|---|---|---|
| **A** | 1 | WT | DMSO | 4 | *zig-1* | DMSO |
| | 2 | WT | WM | 5 | *zig-1* | WM |
| | 3 | WT | DMSO | 6 | WT | WM |
| **B** | 1 | WT | DMSO | 4 | *zig-1* | DMSO |
| | 2 | WT | WM | 5 | *zig-1* | WM |
| | 3 | *zig-1* | DMSO | 6 | HOBO temperature recorder | |
| **C** | 1 | WT | DMSO | 4 | *zig-1* | DMSO |
| | 2 | WT | WM | 5 | *zig-1* | WM |
| | 3 | *zig-1* | WM | 6 | *zig-1* | WM |
| **D** | 1 | WT | DMSO | 4 | *zig-1* | DMSO |
| | 2 | WT | WM | 5 | *zig-1* | WM |
| | 3 | *zig-1* | DMSO | 6 | HOBO temperature recorder | |

**Supplementary Table 2**. Descriptive statistic results for vacuole number assay of cells from epidermis/cortex transition/elongation zone.

| Assay type | | Ground control | | | | Flight assay | | | |
|---|---|---|---|---|---|---|---|---|---|
| Treatment | | WT DMSO | WT WM | *zig-1* DMSO | *zig-1* WM | WT DMSO | WT WM | *zig-1* DMSO | *zig-1* WM |
| Epidermis transition | Mean ± Std. Deviation | 1.2 ± 0.5 | 1.1 ± 0.4 | 2.6 ± 1.1 | 1.4 ± 0.8 | 1.2 ± 0.5 | 1.2 ± 0.6 | 2.6± 1.4 | 1.7 ± 1.0 |
| Epidermis elongation | Mean ± Std. Deviation | 1.0 ± 0.2 | 1.0 ± 0.2 | 3.3 ± 1.5 | 1.7 ± 1.1 | 1.1 ± 0.3 | 1.1 ± 0.3 | 3.8 ± 2.0 | 2.9 ± 1.8 |
| Cortex transition | Mean ± Std. Deviation | 1.2 ± 0.5 | 1.2 ± 0.6 | 2.6 ± 1.0 | 1.5 ± 0.8 | 1.2 ± 0.6 | 1.2 ± 0.5 | 2.6 ± 1.4 | 1.6 ± 1.1 |
| Cortex elongation | Mean ± Std. Deviation | 1.0 ± 0.1 | 1.0 ± 0.1 | 2.5 ± 1.2 | 1.4 ± 0.8 | 1.0 ± 0.3 | 1.0 ± 0.2 | 2.9 ± 2.1 | 1.4 ± 1.0 |

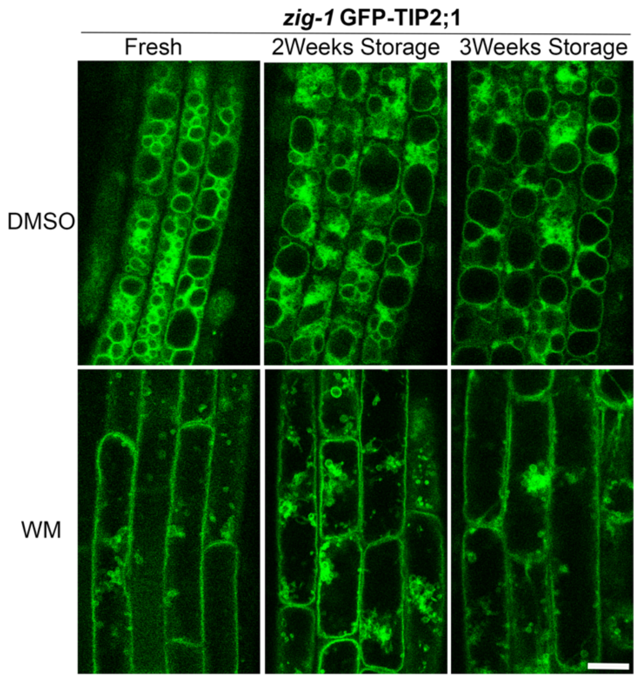


**Supplementary Figure 1**. DMSO and WM solution were still functional after 2-3 weeks storage at 4°C.
*zig-1* seedlings expressing GFP-TIP2;1 were treated with fresh DMSO/WM solution and DMSO/WM that had been stored for 2-3 weeks. 2-3 weeks old DMSO did not change cell physiology while 2-3 weeks old WM were still functional to induce vacuole fusion of *zig-1*.
Scale bar: 20 μm

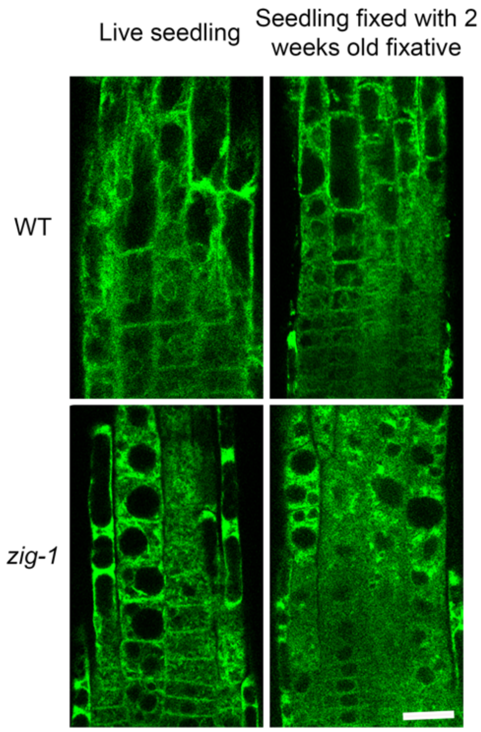


**Supplementary figure 2**. Fixative (4%PFA in 0.5X PBS buffer) was still functional after 2 weeks storage at 4°C.
WT and *zig-1* seedlings expressing GFP-TIP2;1 were fixed by 2 weeks old fixative solution. Those old fixative solution could still preserve cell morphology comparing with live seedling images.
Scale bar: 20 μm

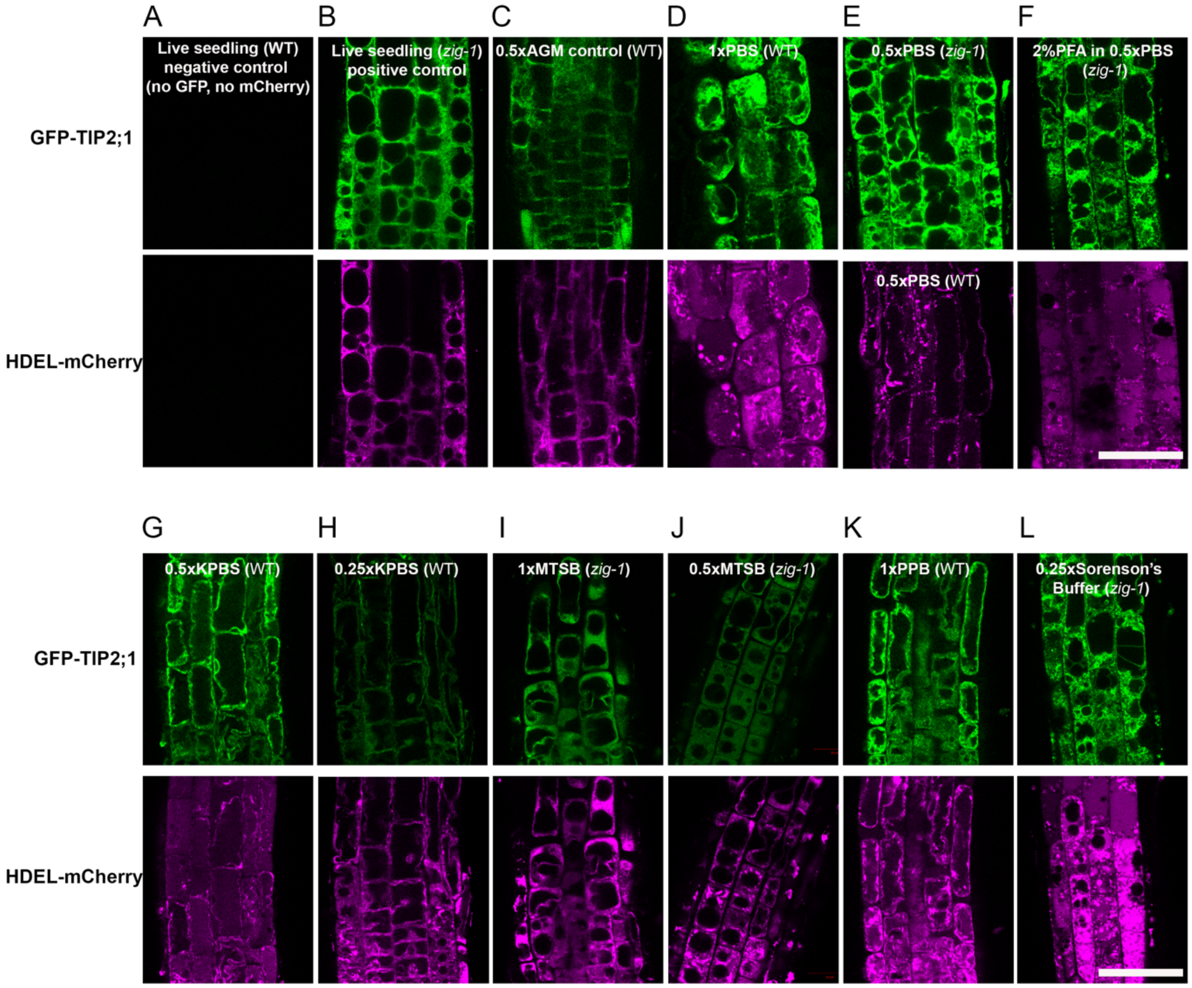


**Supplementary figure 3**. WT (**A, C, D, G, H, K**) or *zig-1* (**B, E, F, I, J, L**) fixed by 4% PFA (except for F) in different buffers.
Three-day-young seedlings grown in dark were put into tubes containing 1 volume of 0.5X AGM for 2 hours in dark. Then another 1 volume of fixative solution were added into tubes to fix seedlings. The final concentration of fixative solution and buffer used are indicated. Confocal images were taken after overnight fixation at 4°C. 4%PFA in 0.5X PBS buffer was the best fixative solution to preserve GFP intensity as well as tonoplast morphology for root tip cells. ER morphology was difficult to preserve in most buffers.
Scale bar: 40 μm

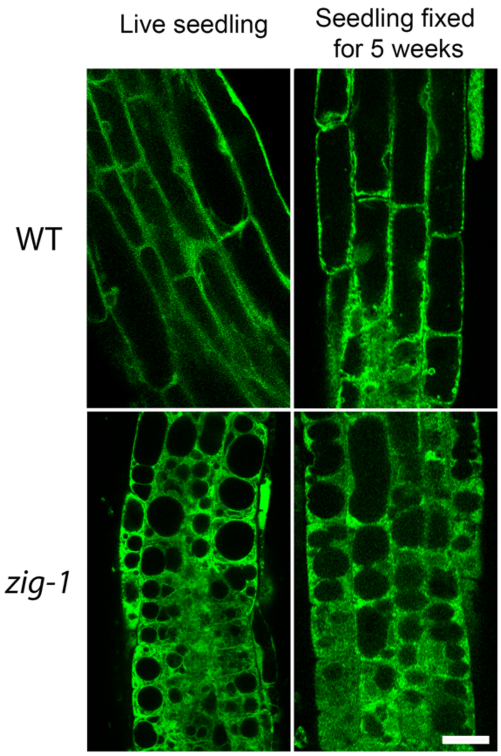


**Supplementary figure 4**. Samples can be stored at 4°C environment for 5 weeks after fixation (fixative is 4%PFA in 0.5X PBS).
Cell morphology and GFP signal of WT and *zig-1* expressing GFP-TIP2;1 can be well preserved as live seedlings after 5 weeks storage in fixative solution.
Scale bar: 20 μm

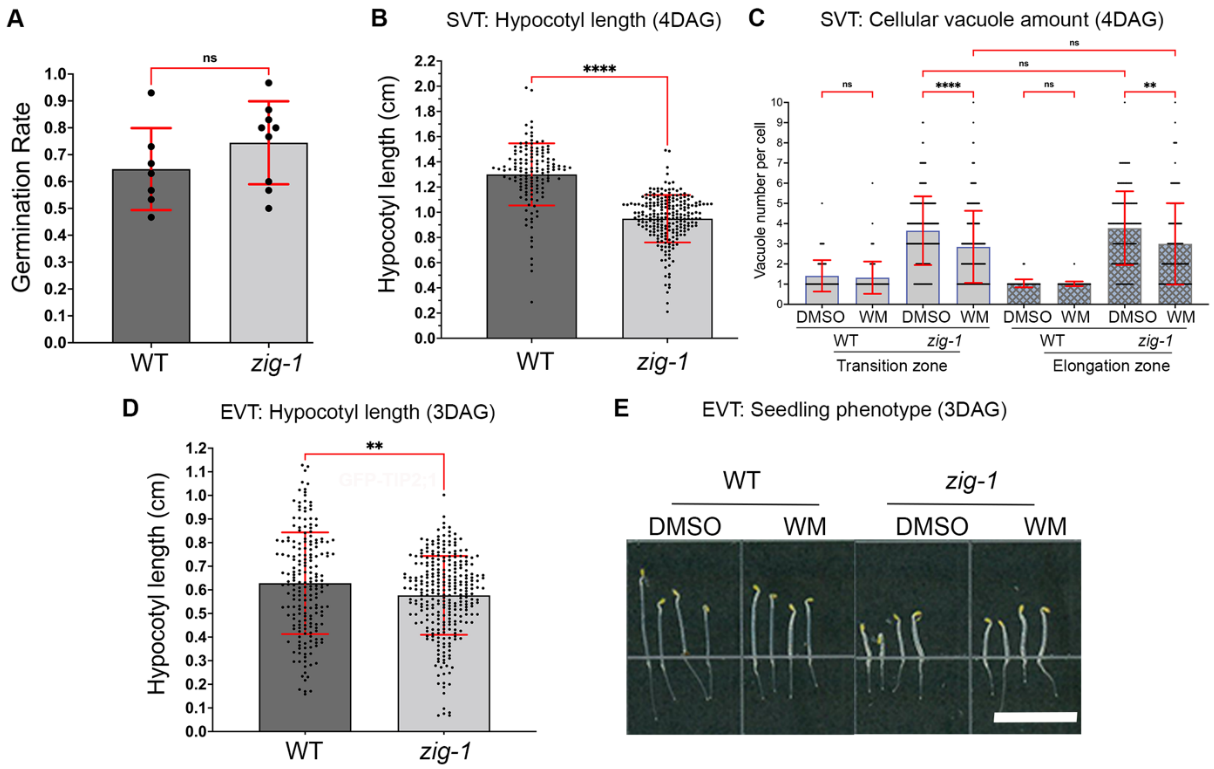


**Supplementary figure 5**. *Arabidopsis* WT and *zig-1* germinated and grew well in BRIC-PDFU from SVT and EVT.

**(A)** Germination rate of WT and *zig-1* grown in the BRIC-PDFU from SVT. No significant differences in germination rate were detected between genotypes (Unpaired t test, ns: p>0.05, n represents plate number, n>=7).

**(B) Hypocotyl length of** WT and *zig-1* grown in the BRIC-PDFU from SVT. WT showed longer hypocotyl than *zig-1* (unpaired t test, **** p<0.0001, n represents seedling number per genotype, n>135). Seedlings were grown at 22°C for 4 days (DAG: days after growth).

**(C)** Cellular vacuole number of WT and *zig-1* from SVT treated with DMSO buffer or WM solution. Cells from both the transition zone and the elongation zone of root tip were analyzed. (3-way ANOVA, Tukey's multiple comparison test, **** p<0.0001; **p<0.0021; n represents cell number under analysis per genotype per treatment, n>68)

**(D)** Hypocotyl length of WT and *zig-1* in the BRIC-PDFU from EVT. Seedlings were grown at 22°C for 3 days. (DAG: days after growth; Unpaired t test, ** p<0.0021, n represents seedling number per genotype, n>196)

**(E)** Representative fixed seedlings of WT and *zig-1* in the BRIC-PDFU from EVT treated with DMSO buffer or WM solution before fixation.

Scale bar: 1 cm

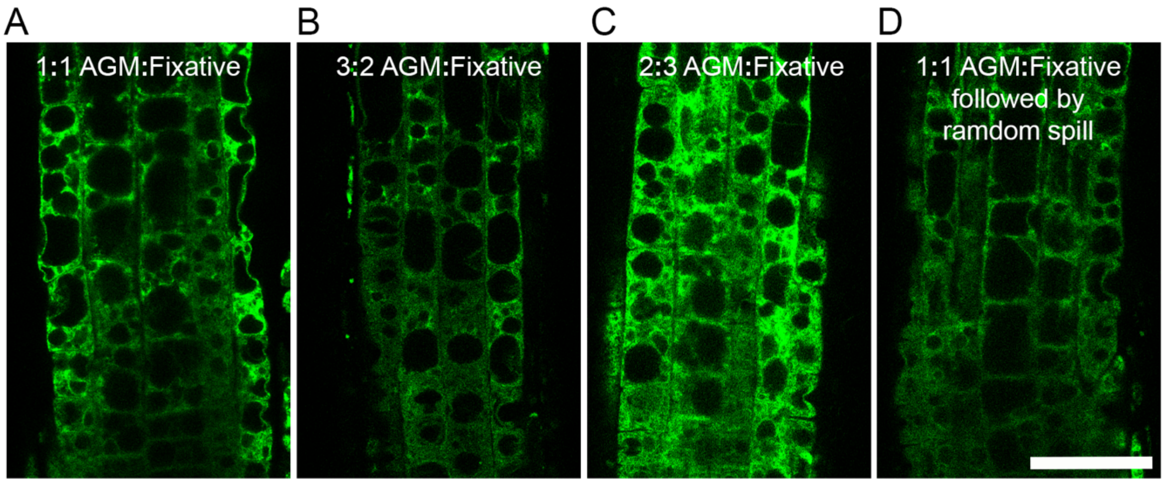


**Supplementary figure 6**. Root tip cells and GFP intensity were well preserved when the AGM and fixative solution were combined with 1:1 ratio.
Root tip confocal imaging for *zig-1* (GFP-TIP2;1) fixed in the AGM and fixative solution (8%PFA in 1X PBS) with a ratio of 1:1 (**A**); 3:2 (**B**); 2:3 (**C**) or 1:1 followed by solution spilling out from plate to mimic the similar liquid loss in the PDFU (**D**).
*Arabidopsis* seedlings were grown in dark for 3 days on petri dishes. Then 0.5X AGM was added to the petri dish for 2 hours in dark. Fixative solution was then added to the petri dishes to fix seedlings. Confocal images were taken after 4°C of overnight fixation.
Scale bar: 40 μm

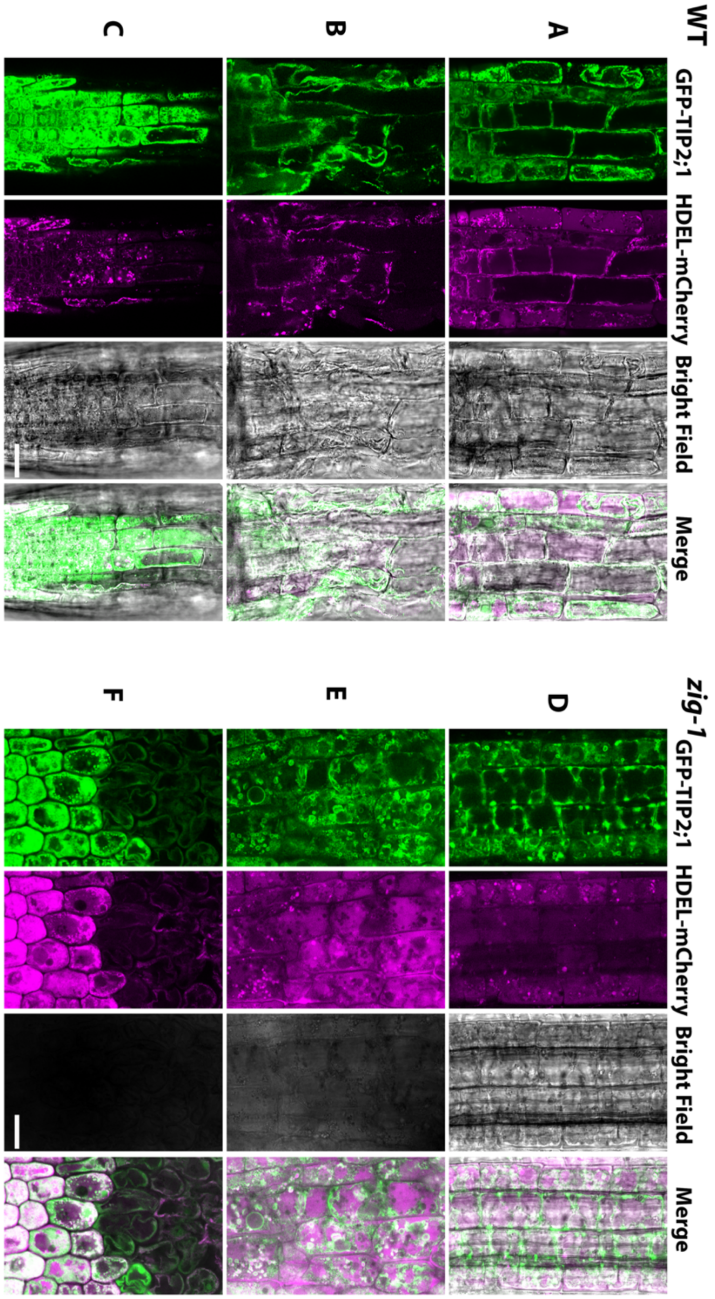


**Supplementary figure 7**. Not all cells from all seedlings can be well preserved during fixation.

For most seedlings, the vacuole in some cells of WT **(A)** and *zig-1* **(D)** can be well preserved, though the ER structure cannot be maintained at the same time. However, there were bad fixation for both WT **(B, C)** and *zig-1* **(E, F)**.

Scale bar: 20 μm

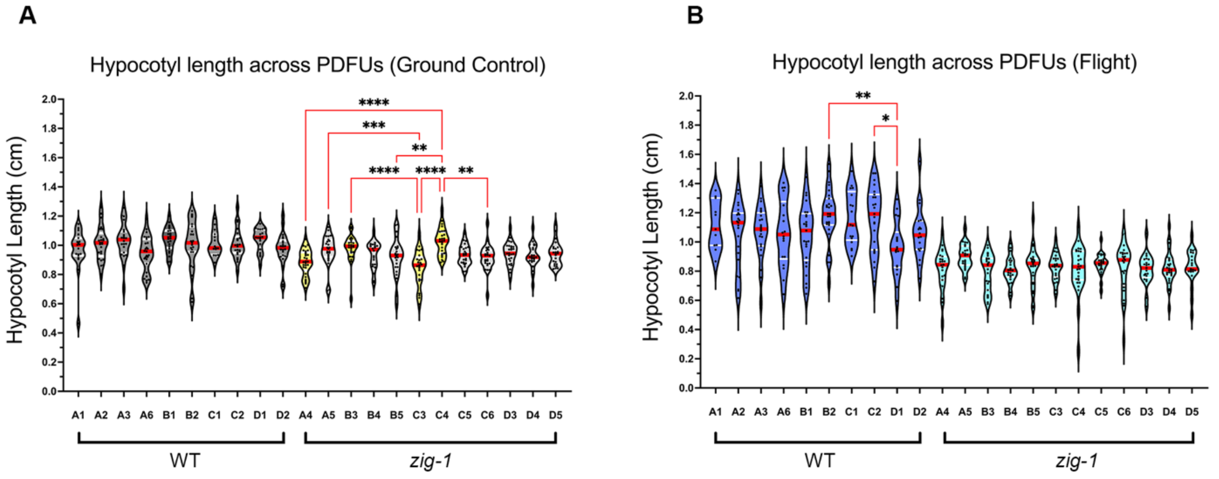


**Supplementary figure 8**. Data of hypocotyl length for WT and *zig-1* from ground control (**A**) and flight assay(**B**) were consistent across different PDFUs.

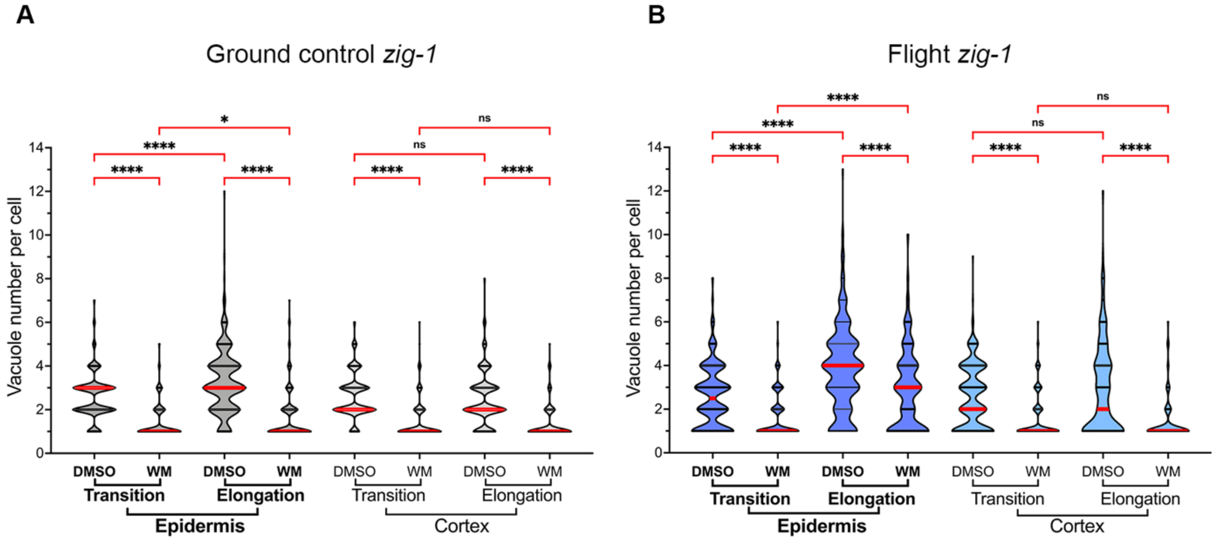


**Supplementary figure 9**. Cells from epidermis transition zone and elongation zone had different cellular vacuole number while cells from cortex transition zone and elongation zone had similar cellular vacuole number considering *zig-1* sample from ground control (**A**) and flight assay(**B**).

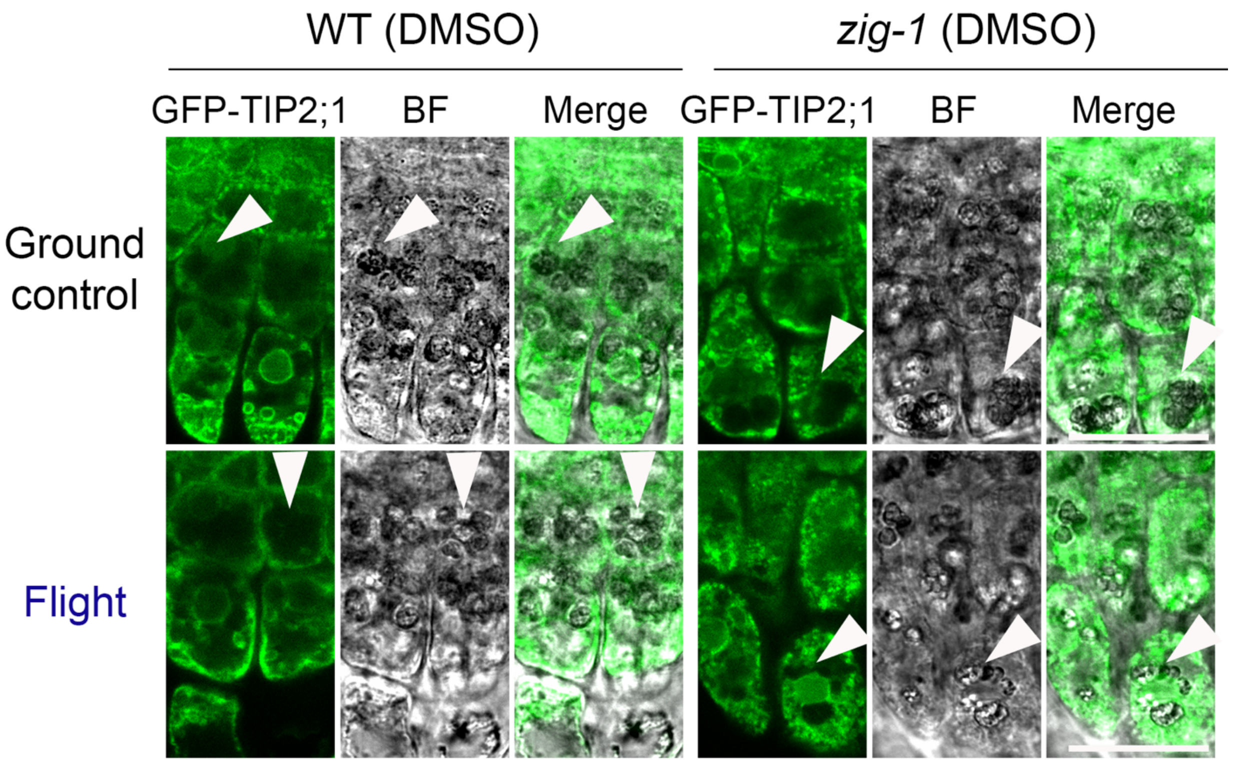


**Supplementary figure 10**. Visualization of amyloplast in columella cells. Amyloplasts (Arrowhead) can be observed in the root tip columella cells for WT and *zig-1* that were grown in the BRIC-PDFU of ground control and flight assay.
Scale bar: 20μm
